# Improving fMRI-based prediction of individual pain ratings by accounting for interindividual variance

**DOI:** 10.1101/2023.07.06.548050

**Authors:** Ole Goltermann, Christian Büchel

**Author notes:** Please address correspondence to: Ole Goltermann, Institute of Systems Neuroscience, University Medical Center Hamburg-Eppendorf, Martinistr. 52, D-20246 Hamburg, Germany.

## Abstract

We challenge the pessimistic conclusion of a recently published paper by Hoeppli et al that fMRI-BOLD measures are useless in predicting interindividual differences in pain perception. By conducting a re-analysis of publicly available data of their study, we propose an alternative analysis approach that addresses the issue of interindividual variance differences in BOLD readouts, aiming to enhance the predictive power of fMRI-based measures. Instead of using absolute values of both, pain ratings and BOLD measures, we make use of robust intraindividual differences between the two experimental conditions in their study. Our findings demonstrate a statistically significant positive linear relationship between the neurologic pain signature (NPS) score, a multivariate pain classifier based on BOLD fMRI, and individual differences in perceived pain ratings. This relationship is driven by individuals that report pain sensitivity to both experimental conditions and can clearly distinguish between the two. Our results provide evidence for the potential of fMRI-BOLD measures in predicting interindividual differences in pain perception and allow for a more optimistic conclusion regarding the ongoing debate whether fMRI can be used as an objective measure for pain perception.

**ARISING FROM:** Hoeppli, M.E., Nahman-Averbuch, H., Hinkle, W.A. et al. Dissociation between individual differences in self-reported pain intensity and underlying fMRI brain activation. Nat Commun 13, 3569 (2022). https://doi.org/10.1038/s41467-022-31039-3

---

In a recent study, Hoeppli et al.^1^ tested whether interindividual differences in the rating of a noxious stimuli (48°C) could be predicted by individual brain activity measured with blood-oxygenated-level dependent (BOLD) fMRI. While they observed that BOLD was sensitive to small differences in applied noxious stimuli within an individual, their analysis revealed no relationship between interindividual BOLD measures and pain ratings. Based on these results, they concluded that fMRI may not be a useful objective measure to infer reported pain intensity. In the following, we aim to challenge this conclusion by re-analysing their data with a focus on reducing interindividual variance. Our results allow for a more optimistic conclusion, demonstrating evidence for predicting interindividual differences in pain perception.

One possible explanation for the absence of a relationship between individual pain rating and the corresponding BOLD readout is the between-person variability of BOLD activity, which has been explored in previous studies^2–4^. To address this issue, several solutions have been proposed. Many of them emphasize the selection of the fMRI contrast, highlighting that BOLD, as a non-quantitative contrast, may not be suitable for predicting individual differences. As promising alternatives, measures for cerebral blood flow or cerebral blood volume changes might reduce between person variability. Moreover, the BOLD signal could be measured in a quantitative manner^5^, calibrated manner^6^ or could be rescaled post-hoc using resting-state data or vascular reactivity maps^7^.

As another way to reduce between-person variance, without the need of using a different fMRI contrast or rescaling the recorded BOLD data, we propose utilizing robust intraindividual differences between experimental conditions. Hoeppli et al. demonstrated that high (48°C) and low (47°C) stimulus intensities reliably differed within individuals in terms of perceived pain intensity, region-specific BOLD activation, and neurologic pain signature (NPS) scores^8^. We hypothesized that by using relative differences between these two conditions instead of absolute values^9^, one can “calibrate” BOLD-based readouts within individuals and thus account for interindividual variance. To test this idea, we reanalyzed the publicly available NPS data from the study conducted by Hoeppli et al.

We calculated a difference score for NPS and perceived pain ratings by subtracting the NPS score and perceived pain rating for high and low stimulus intensities, respectively^10^. We then entered the individual NPS differences for all subjects (N = 101) into a linear regression model to predict individual differences in perceived pain ratings. Our findings revealed a small yet statistically significant positive linear relationship (*r* =.20, *p* =.048, see Figure 1a). It is important to note that this approach might have yielded an even more robust relationship when using the raw BOLD data, which unfortunately was not made publicly available. In addition, the heat task included only four low intensity noxious stimulus (47 °C), but 17 high intensity noxious stimuli (48 °C). It is to be expected that a more balanced design would have further improved the predictive value of BOLD responses.

**Figure 1.**
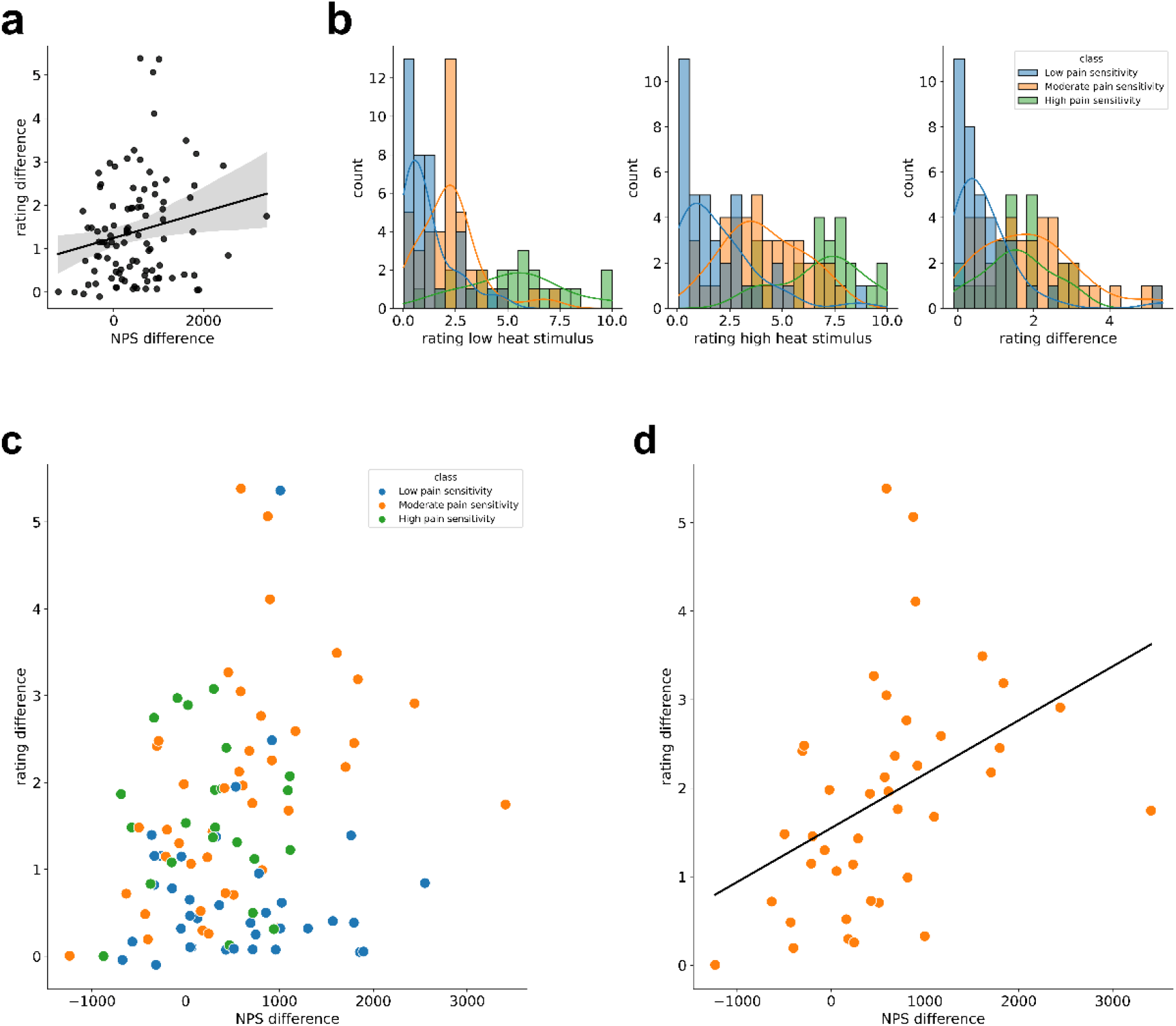
**(a)** Predicting differences in ratings for high vs. low intensity stimuli using differences in NPS between high vs. low intensity stimuli, *r* =.20, *p* =.04 **(b)** Histograms for ratings of low and high intensity stimuli, as well as the difference between the two, color-coded for the three subgroups of low, moderate and high pain sensitivity **(c)** Relationship between difference in perceived pain ratings (high vs. low) and difference in NPS scores (high vs. low), color-coded for the three subgroups of low, moderate and high pain sensitivity **(d)** Predicting differences in ratings for high vs. low intensity stimuli using differences in NPS between high vs. low intensity stimuli for moderate pain sensitivity group, *r* =.42, *p =*.*0063*)

Next, we examined the distribution of perceived pain ratings for low and high intensity stimuli, as well as the difference between the two. For this we split the data into the three subgroups of low, moderate and high pain sensitivity subjects, according to the behavioural data of Hoeppli et al. (defined with data *prior* to fMRI data collection). Interestingly, a considerable number of subjects (*n* = 29) reported a perceived pain below two (on a scale from zero to ten), even for the high intensity heat stimulus. It is worth noting that for these subjects, the predictability of pain ratings by NPS scores is questionable, given their minimal or absent pain experiences. Furthermore, the distributions of ratings for the low, high, and difference scores are skewed to the right in the low pain sensitivity group (Figure 1b). To investigate the relationship between NPS differences and rating differences in more detail, we plotted the data color-coded for the three subgroups of low, moderate, and high pain sensitivity (Figure 1c). We conducted linear regression analyses using the individual NPS differences for each subgroup to predict individual differences in perceived pain ratings. No statistically significant relationship was found between NPS differences and rating differences for the low (r =.07, p =.69, n = 37) and high (r = -.04, p =.84, n = 23) pain sensitivity groups. However, a statistically significant linear relationship was observed between NPS differences and rating differences for the moderate pain sensitivity subgroup (r =.42, p =.0063, n = 41, Figure 1d).

Based on our results, we draw three conclusions. First, by utilizing relative differences instead of absolute values, we provide modest evidence for predicting interindividual differences in perceived pain ratings using fMRI-BOLD based NPS scores (Figure 1a). Second, we establish a medium-sized correlation between differences in NPS and differences in perceived pain ratings for the moderate pain sensitivity group (Figure 1d), supporting our first conclusion and suggesting that the success of prediction may be influenced by the distribution of the dependent variable (pain ratings) within the sample. Finally, considering that a significant proportion of subjects reported low or no pain even in response to the high intensity stimulus, we suggest that prediction accuracy may improve by using relative differences between stimuli, where one clearly evokes a painful experience and the other is substantially lower. This implies that the choice of stimuli used in Hoeppli et al. (47°C and 48°C) may be suboptimal for predicting interindividual differences and could therefore underestimate predictive power. Future studies should explore whether selecting more distinct stimulus intensities can enhance prediction success. We further recommend re-analysing existing fMRI data with a focus on robust relative intraindividual differences between experimental conditions when predicting interindividual differences. Additionally, exploring the potential benefits of post-hoc rescaling of BOLD data using resting-state or vascular residuals to reduce interindividual variance would be a valuable direction for future investigation.

